# Decoding human lifespan from neural noise and explaining age-related changes in fractal dimension and gamma oscillations using fractional harmonic oscillator

**DOI:** 10.1101/2025.09.13.675905

**Authors:** Srishty Aggarwal

## Abstract

Predicting human lifespan is a longstanding objective of biomedical research. Traditional statistical models estimate mortality risk or biological age but not lifespan. We propose a two-parameter model based on stochastic fractional harmonic oscillator for neural signals. The model computes maximum human lifespan using 1/f slope, the measure of neural power decay with frequency, which is indicative of neural noise. Using slope rate from electroencephalographic and electrocorticographic datasets, we estimate the mean lifetimes of healthy adults and epileptic patients as 76.9 and 69.7 years, resulting in 89.4% and 96.9% accuracy respectively. Additionally, the present model resolves the inconsistency in age-related changes of fractal dimension (FD) and captures naturally the non-monotonic variation of stimulus-induced gamma power. Thus, the present model provides a simple way to estimate lifespan while explaining age-related changes in slope, FD and power simultaneously, thereby, paving the way for individualized lifetime measurements and unveiling fundamental principles governing life.

**One line Summary:** We build a two-parameter model for neural signals that predicts mean human lifespan through a single neural observable, that may lead to future individualized lifetime predictions along with unveiling fundamental principles of life.

## Introduction

Predicting human lifespan is a longstanding goal of biomedical research, with implications spanning healthcare, policy, and the understanding of biological ageing. Existing models rely on demographic data (*1*), molecular markers such as DNA methylation (*2*), composite physiological indices including blood pressure (*3*) and heart rate variability (*4*), and machine learning model like ‘life2vec’ (*5*). However, these approaches mainly focus on mortality risk after a certain age or computation of present biological age, not on actual lifespan. They depend on statistical approaches with multiple parameters and therefore, vulnerable to data-specific biases. This limits their ability to get insights about the fundamental processes underlying ageing, disease and lifespan that can be applied universally. Hence, there is a need for pure theoretical model that can predict lifespan using a single feature.

Neural signals, derived from electroencephalography (EEG) and electrocorticography (ECoG), exhibit reproducible, individual-specific patterns (*6*) and systematic changes across the lifespan (*7–10*). In particular, the slope of the neural power spectral density (PSD), a measure of how signal power decays with frequency, shows a robust, age-dependent decline from infancy to old age (*7, 10–12*). This “1/f” slope, often known as neural noise, has been linked to circuit-level characteristics such as synaptic filtering (*13, 14*) and excitation-inhibition balance (*15*), as well as arousal level (*16*) and task performance (*17*). Another PSD feature, the peak representing the oscillatory activity in different frequency range like alpha (8-12 Hz) and gamma (20-66 Hz) also exhibit age-related variation (*18, 19*). The peak power of gamma oscillations, elicited by visual stimulus (*20*) modulated by attention and cognitive memory (*21*), varies non-monotonically with age, while increasing from childhood to adolescence and decreasing later (*8, 19*).

While these neural features show promise as physiological markers of ageing, researchers could not capture their age-related variation together using a single model (*22*). Most models address the dynamics governing only one feature at a time (either power or slope) (*23–26*) and assume memoryless or Markovian dynamics, i.e., neural activity depends only on its current state. However, the brain is a non-linear system with memory and fractal-like properties that span multiple timescales (*27, 28*) and, thus, requires a model that could deal with such properties. Moreover, signal’s non-linearity, often measured by Higuchi fractal dimension (HFD), exhibits inconsistent changes with ageing (*9, 29, 30*) that awaits a theoretical resolution.

Therefore, to capture the non-Markovian and fractal nature of neural signals, we model them via stochastic fractional harmonic oscillator (FHO). It intrinsically includes memory through its non-integer derivatives. We use this framework to estimate human life expectancy in existing EEG and ECoG datasets through slope rate. We further validate this model by providing a means of resolving the conflicting findings of HFD with by understanding its underlying non-linear properties, and explaining age-related changes in power and centre frequency of stimulus-induced gamma oscillations.

## Results

### Model

We consider that the neural signal *X(t*) is governed by the stochastic FHO, described as

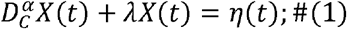

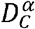 is the Caputo fractional derivative of order *α* with 1 < *α* 2, *λ* is the decay constant, lying between 0 and 1, and *η* is the random Gaussian noise. The Caputo derivative for *α* between 1 and 2 is represented as convolution of a ‘memory term (1/*t* ^*α* -1^)’ with the second order derivative of *X* (*t*). As memory depends on *α* only, *α* is the main representative of non-Markovian behaviour in FHO, such that with decreasing *α*, the present behaviour of a system depends more on its past states, i.e., the system retains more memory. To distinguish this memory from the cognitive or neural memory, we call it ‘chronomemory’ (the memory about the history of the system accumulated over time).

First, we observe the variation of the time series *X*(*t*) and its features with *α* and *λ*. It forms the basis for our assumptions of various constraints on *α* to explain neural signals. For this, we solve Eq. (1) to obtain *X* (*t*) We extract the signal features, namely, 1/f slope and centre frequency (CF) and peak height (PH) of oscillations from PSD and fractal dimension (FD) analytically as well as numerically (see Materials and Methods).

Figure 1A illustrates the time series for a single trial (repeat) and the corresponding trial-averaged PSDs, obtained numerically, at different *α* for *λ*= 0.3 (top) and 0.8 (bottom). At *λ* = 0.3, time series at low *α* (1.2 and 1.45) have low amplitude and appear noise-like (Fig. 1A top left). As *α* increases, they start displaying oscillatory behaviour. This is also observed in the corresponding PSDs (Fig. 1A top right), where low *α* are characterized by bending (knee), and pronounced peaks are present only for higher *α* (1.7 and 1.95). The PSD slope, computed in the frequency range of 64-140 Hz, becomes sharper (faster power decay) as *α* increases. Further, the numerical FD (here, HFD) decreases with increase in *α* as indicated in legends. The same trend could be seen for *λ* = 0.8 in Fig. 1A bottom panels, with the peaks shifting towards greater frequencies as compared to *λ* = 0.3 in PSDs. Figure 1B represents results analogous to Fig. 1A but for different *λ* at *α* = 1.15 (top) and 1.9 (bottom). The distinction between the shape of PSDs at low vs high *α* is clearly evident in the top and bottom panels. At low *α* (1.15), the time series are noisy with no oscillatory peaks for all *λ* (Fig. 1B top). The corresponding PSD slope is also similar for all *λ*. In contrast, oscillations emerge at high *α* (1.9), and *λ* mainly plays a role in modulating PH and CF of these oscillations (Fig. 1B bottom). Though the PSDs for these *λ* at high frequencies seem to merge, the slope is slightly higher for a larger *α*, due to larger power at low frequency end (∼50 Hz), resulting in a faster decay for greater *λ*. HFD increases slowly with *λ* at both *α*.

**Figure 1.**
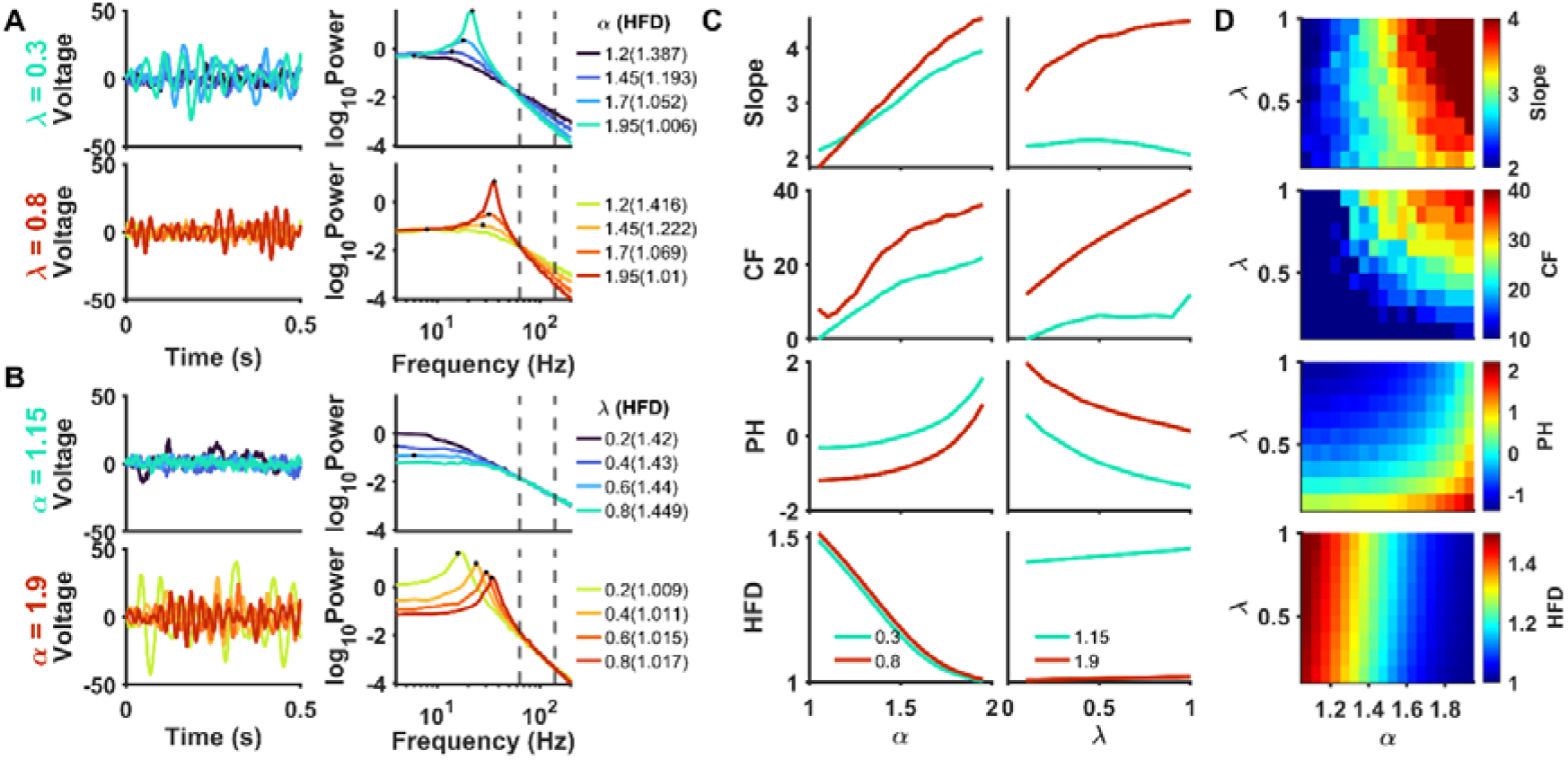
Variation of signal features with *α* and *λ* for stochastic fractional harmonic oscillator (FHO). **(A)** Time series (left) and trial – averaged power spectral densities (PSDs; right; averaged for 200 trials) for *α* 1.2, 1.45, 1.7 and 1.95 for *λ* = 0.3 (top) and 0.8 (bottom). The knee/ peak, for which centre frequency (CF) and peak height (PH) is computed, is highlighted by a black dot for each PSD. The slope is computed in the frequency range of 64-140 Hz, indicated by grey lines. The Higuchi fractal dimension (HFD) for the time series is mentioned in the legend along with the corresponding *α*. **(B)** Same as (A) but at different *λ* (0.2, 0.4, 0.6 and 0.8) for *α* = 1.15 (top) and 1.9 (bottom). **(C)** The variation of slope, CF, PH and HFD with *α* (left) and *λ* (right) at *λ* and *α* chosen in (A) and (B) respectively. **(D)** The simultaneous variation of these features with *α* and *λ*.

The associated variation of numerically computed slope, CF, PH and HFD with *α* and *λ* individually and combined is summarized in Fig. 1C and D respectively, that can be compared with analytical relations. Analytically, the slope increases with increase in with maximum value as 2*α* while it decreases with increase in *λ*. The numerical slope increases almost linearly with *α* (Fig. 1C top left panel) but crosses ′2*α*′ limit at high *α* for *λ* = 0.8. Further, it decreases slowly with increase in *λ* at *α* = 1.15 following analytics but exhibited opposite trend at high *α* (Fig. 1C top right panel) that is inconsistent with the analytical results. These inconsistencies of slope variation at high *α* are due to distortion of slope computation in presence of oscillations (*10, 31*). However, the peak characteristics are not distorted due to slope and show same behaviour as analytics with the CF of knee (at low *α*)/ peak (at high *α*) increasing with both *α* and *λ*, and the PH increasing with *α* and decreasing with *λ*. There is a steep increase in PH at *α*> 1.6, due to emergence of oscillatory peaks. Further, HFD decreases with increase in *α*, following the theoretical relation between *α* and FD, but it also exhibits a weak dependence on *λ*.

Since oscillatory peaks predominantly appears only at high *α* and slopes get affected in presence of oscillations ((*10, 31*) and (supplementary section 1)), going forward, we constrain *α* between 1 and 1.5 for the slope computation and 1.5 to 2 for the peak features such as CF and PH. The separate *α* range for slope and oscillations would enable us to capture their dynamics independently.

### Assumptions to explain neural features

To explain the age-related changes in neural signals, we have made two assumptions:

1. *α lies between 1 and 1.5 for baseline and 1.5 and 2 for stimulus-induced activity*.
2. *α in baseline range decreases ‘uniformly’ with age from 1.5 at birth to 1 at death*.

The first assumption is based on the general characterization of baseline activity using slope (*10*) and (visual) stimulus activity by gamma oscillations (*8*). The transition at 1.5 provides equal *α* range for baseline and stimulus. Further, as baseline slope was shown to decrease with age from infancy to old age (*7, 10–12*) with linear fitting at different age stages, we assume that slope decreases at a constant rate from birth till death. As slope varies linearly with *α* (Fig. 1(C)), it leads to the second assumption (Supplementary Section 4.1).

### Human life expectancy using baseline slope

We try to estimate human lifespan using the assumption 2., i.e., the uniform change of baseline *α* from 1.5 at birth to 1 at death. We use slope as a measurable parameter for the estimate of human lifetime, as it shows the monotonic decrease with age (*7, 10–12*). Thus,

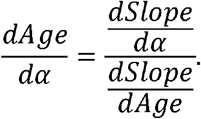

*dSlope/dα* is less than or equal to 2 (Eq. (12), Materials and Methods) and *dSlope/dAge* or the ‘slope rate’ can be measured from the data. Therefore,

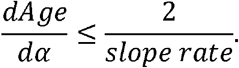

Therefore,

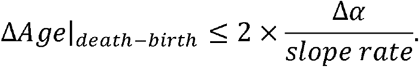

As Δ*α* = -0.5 from birth to death (as per assumption 2),

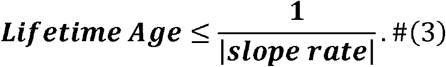

The above expression turns into equality when *dSlope/dα* = 2. This could be attained with an appropriate choice of frequency range for slope computation as shown in Fig. S1.

### Computation of age using neural data

We take two datasets for the estimation of life expectancy, an EEG dataset of healthy individuals from (*10*) and an ECoG dataset of epileptic patients from (*7*). The details of these datasets are mentioned briefly in Material and Methods.

#### Healthy Individuals

The EEG dataset consists of eyes closed data from 204 healthy subjects aged 50-88 years from Bengaluru, Karnataka, recorded under Tata Longitudinal Study of Aging (TLSA) (*10*). In the frequency range of 64-140 Hz, the slope rate is − 0.013 ± 0.0004 (mean ± standard error of mean (SEM); p = 4.77 × 10^−7^, one-tailed test, *t-statistics*) for a group of parietal and occipital electrodes (*10*), as also depicted in Fig. 2A. This slope rate results in maximum lifetime age of 76.9 [74.6, 79.4] years using Eq. (3) (the bounds on the age are obtained by using the inverse of ‘slope rate SEM’). It aligns with 89.4% accuracy with the regional life expectancy at birth data (69.5 years in Karnataka, India between 2015-2019, (*32*)). Since the separate data for human life expectancy for healthy individuals are not available, we compare with the mean life expectancy at birth data of the regional population for the years around which the present data has been recorded. The life expectancy for individual gender and its dependence on electrode position is discussed in Supplementary Section 2.

**Figure 2.**
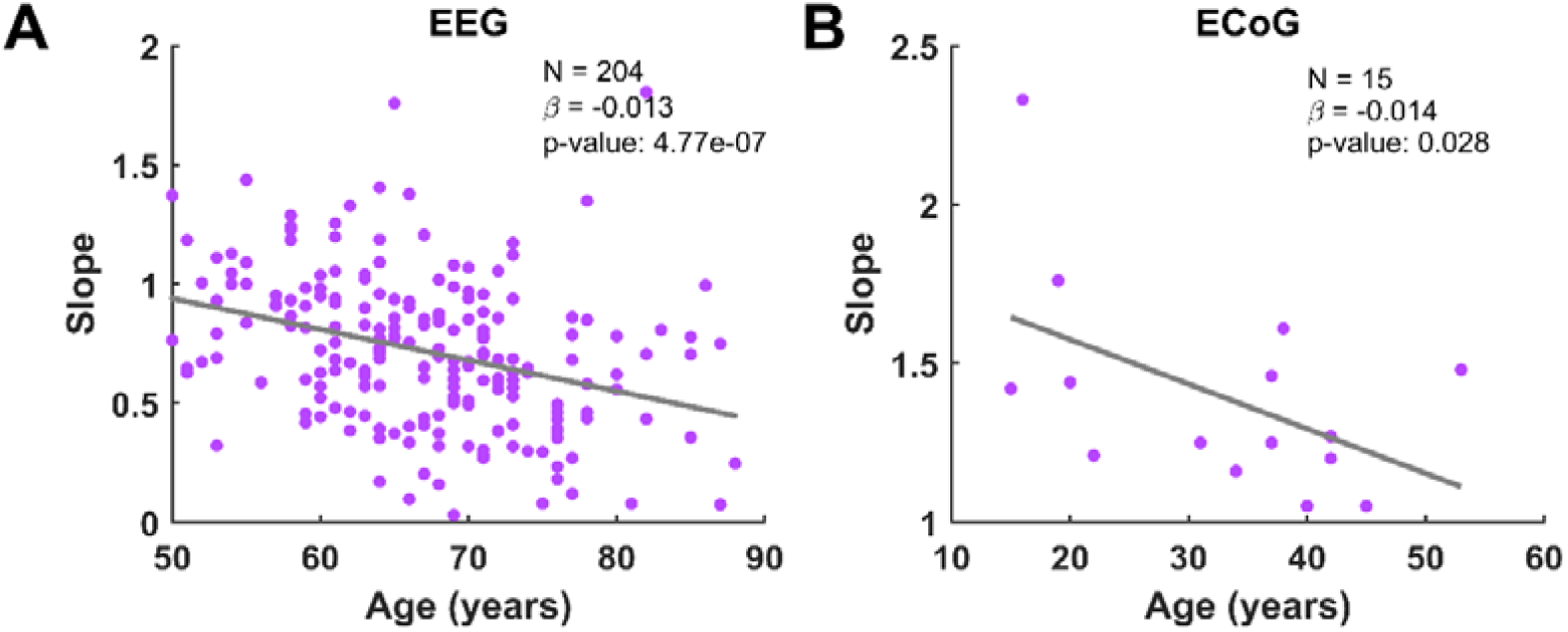
Scatter plots showing the variation of slope with age. **(A)** The EEG eyes closed data (Figure adapted from Fig. 6 of (*10*)). The slopes are computed in frequency range 64-140 Hz. **(B)** The ECoG data (Figure adapted from Fig. 2B of (*7*)). The slopes are computed in frequency range 75-150 Hz. The regression line is in grey. The number of subjects (N), the regression slope (*β*) and the p-value using one tailed test, *t-statistics* are indicated in each subplot.

#### Epileptic patients

The ECoG dataset is constituted of 15 epileptic patients aged 15-60 years, recorded from San Francisco (*7*). The data were analysed from frontal, temporal and supramarginal neocortical regions. For these patients, the slope rate is − 0.014 ± 0.004 (*p* = 0.028, one-tailed test, *t-statistics*; Fig. 2B) in the frequency range of 75-150 Hz, giving maximum lifetime age of 69.7 [55.5, 100] years. The mean life expectancy in San Francisco in year 2011-2015 is 82.9 years (*33*), 13.2 years higher than the calculated maximum life expectancy of 69.7 years for epileptic patients. Interestingly, this difference is close to the mortality rate difference of ∼11 years between healthy cohort and epileptic patients found in Danish population by Drier and colleagues (*34*). Therefore, the present lifespan estimation for epileptic patients is 96.9% accurate, following (*34*).

Thus, slope rate gives a practical lifetime expectancy that lies close to the observed mean human lifespan between 60-100 years. The datasets chosen for present age estimation gives good approximation due to slope computation in high frequency range that matches with our model predictions range (with centre frequency > 50 Hz; Fig. S1 and supplementary section 4.2). This is in contrast with most EEG studies which computed slopes in frequency range ∼1-40 Hz (*11, 12, 18, 35*), thus, making them unsuitable for lifespan estimation. Further, the precision (the average deviation from the mean) in slope for the present EEG and ECoG datasets are 0.25 (34.8%) and 0.23 (16.5%) respectively. It implies that for reliable slope rate estimation from a longitudinal data for individualized lifespan estimation with mean slope rate ∼ − 0.013, one needs to record with ∼20 years gap between the two recordings of an individual. Future techniques for better slope precision could help with faster and reliable slope rate measurements.

#### Explanation of age-related changes

To assess the suitability of our model as a descriptor of neural dynamics, we use it to explore the age-related changes in neural signal features: HFD and PH and CF of gamma oscillations, during baseline and stimulus as shown below. We also compare our model with the damped harmonic oscillator (DHO) based auto regressive (AR2) model (supplementary section 3) which fails in explaining these concomitant results, thus, demonstrating the superiority of our approach.

### Baseline HFD variation with age and slope

There had been an inconsistent variation of HFD with age in literature (*9, 29, 30*). While some studies reported a non-monotonic pattern, with HFD increasing till around 50 years and decline thereafter (*29, 30*), Aggarwal and Ray (*9*) observed an increase in HFD between 50 and 88 years. To address this discrepancy, we utilize the present model that systematically characterizes the age dependence of HFD.

We examine the variation of HFD in baseline by analyzing the signals with *α* between 1 and 1.5, based on assumption 1. The resting state or baseline activity at different ages is represented by triangles in *α* − *λ* − HFD plot (Fig. 3A top panel). The triangles are uniformly spaced (by assumption 2) and become lighter to darker with increasing age. The decreasing *α* led to increasing baseline HFD with age, thus, providing an upper hand to the results reported by Aggarwal and Ray (*9*), that HFD increases with age, over the studies that showed the opposite (*29, 30*). Further, the slope in the frequency range 64-140 Hz decreased uniformly with ageing, (Fig.3B top and bottom panels; *β* = −0.0121; *p* = 3.13 × 10^−4^, *t*−*test*), in consistency with the earlier observational studies (*7, 10–12*). Moreover, HFD and slope followed an inverse relation (Fig. 3A bottom panel; correlation coefficient *ρ* = − 0.99, *p* = 2.67 × 10^−4^), similar to the strong negative correlation (-0.99) between age-related changes in baseline HFD and slope as shown by Aggarwal and Ray (*9*). These results indicate that our present model effectively reconciles the divergent findings of HFD with ageing, along with satisfying the negative correlation between HFD and slope.

**Figure 3.**
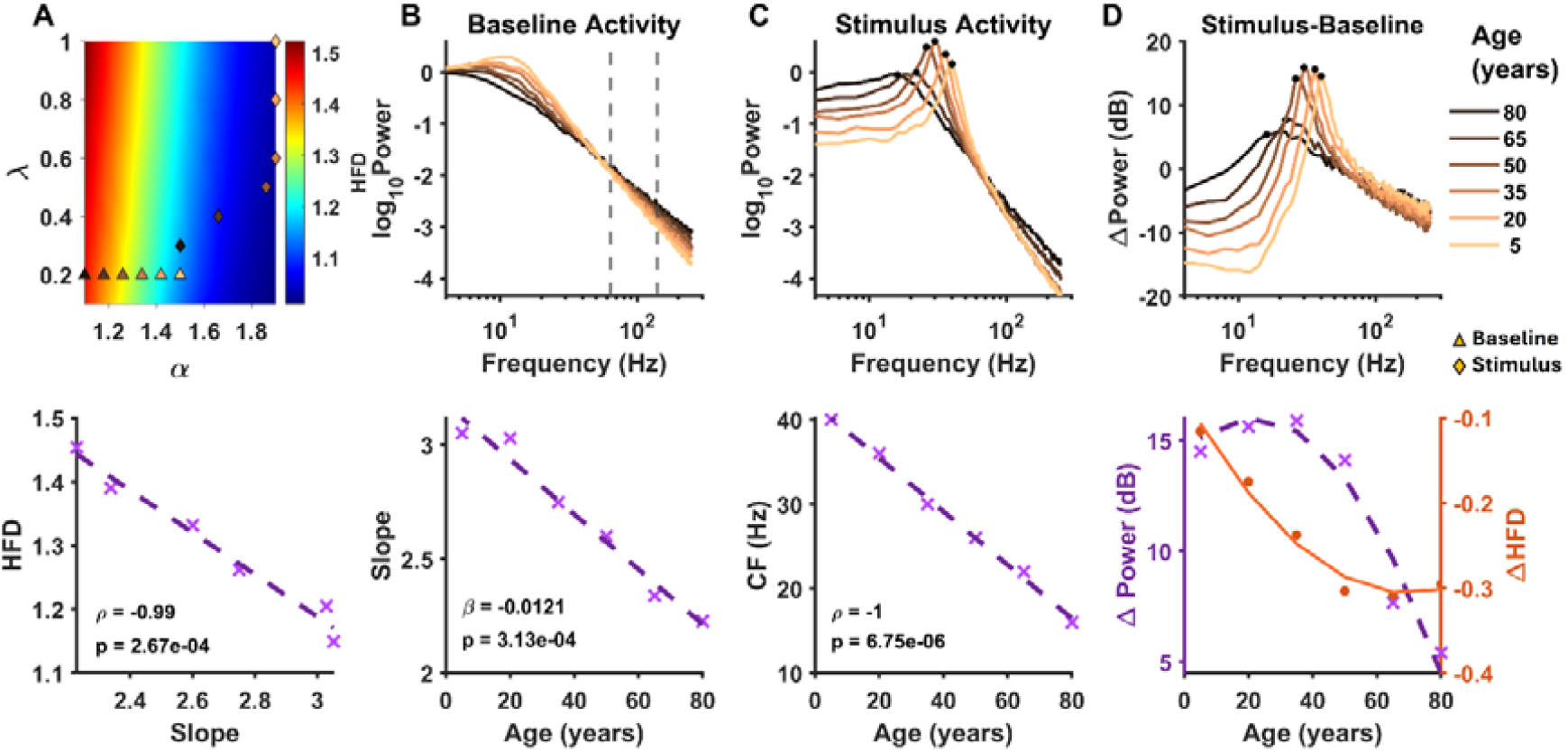
Examination of age-related changes in baseline and stimulus activity. **(A)** (Top) The colormap showing HFD variation with and (HFD plot). The triangles and diamonds represent the baseline and stimulus activity respectively. The shades from lighter to darker tone represent ageing from younger to older age. (Bottom) The variation of baseline HFD with slope computed in frequency range of 64-140 Hz. ‘x’ represents the HFD and slope value corresponding to a triangle in the HFD plot. The dashed line represents the regression curve. The correlation coefficient () and the p-value using t-test are mentioned. **(B)** The variation of baseline PSDs (top) and the slopes (bottom) with age corresponding to the triangles in HFD plot. The regression slope) and the p-value using t-statistics are indicated in the bottom plot. **(C)** (Top) The PSDs corresponding to the diamonds representing the stimulus activity for different ages in HFD plot. The oscillatory peaks are marked by black dots. (Bottom) The variation of corresponding CF with age. and p-value are mentioned. **(D)** The variation of change in power with age (Power in dB; obtained by the difference of log_10_ Power of stimulus (shown in (C)) and baseline (shown in (B)) and multiplying it by 10). The black dots represent the Power at CF marked in (C) top panel. The corresponding Power is plotted as ‘x’ in the bottom panel along with the regression curve. The orange dots and the regression curve in the bottom panel represent HFD, which is obtained by subtracting the HFD at diamond from the triangle for each age group.

### Stimulus-induced oscillations and ΔHFD

We plot diamonds in Fig. 3A top panel to represent the stimulus-induced activity. Based on assumption 1, they are confined between 1.5 and 2 and follow the same age-related color scheme as the baseline. Previous studies have shown an age-related decline in CF of gamma oscillations induced by visual grating stimulus (*8, 19*). As CF in our model is *λ* dependent (Fig. 1B bottom panel), we choose the decrement of *λ* with age for stimulus activity (Fig. 3A top panel). It results in the uniform age-related decrease of CF of stimulus activity, as shown in the stimulus-induced PSDs in Fig. 3C top panel and its variation with age (*ρ*= -1; p =6.75 ×10^−6^, t-test) in Fig. 3C bottom panel.

Aggarwal and Ray (*9*) reported a decline in HFD due to visual grating stimulus i.e., ΔHFD (stimulus — baseline) was negative; with more decline in mid group (50-65 years) than the old one (65-88 years). Since, *α* range for baseline (1-1.5) is lower than for stimulus (1.5-2), it results in higher HFD in baseline than stimulus, leading to negative ΔHFD, thus, consistent with the observed results. Further, as HFD varies almost linearly with *α* (Fig. 1C bottom left panel), ΔHFD∼ *c*Δ *α*, where ‘c’ is a constant. The larger negative ΔHFD in the mid-age group compared to the old-age group (*9*) (|ΔHFD|_*mid*_ > |ΔHFD|_*old*_ ; taking absolute values) implies that |Δ *α* |_*mid*_> |Δ *α* |_*old*_ i.e., Δ *α* increases with decrease in age. For the oldest age group with baseline *α* around 1, the minimum possible Δ *α* can be 0.5 (taking *α* _*stimulus*_∼1.5). Therefore, |Δ|_*mid*_ must be greater than 0.5. However, for a child with *α* _*baseline*_ around 1.5, *α* _*stimulus*_ can maximally be 2, resulting in maximum |Δ *α* |_*child*_ to be 0.5. In summary,

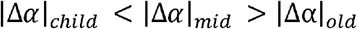

Thus, Δ*α* and thereby, ΔHFD follows a non-monotonic behaviour with age as shown in Fig. 3D bottom panel. It results in the non-monotonic change in the peak height of ΔPower (in dB; obtained by subtracting the log power of baseline from that of stimulus and multiplying by 10) with age as depicted in Fig. 3D top and bottom panels. This non-monotonic behaviour of stimulus induced gamma power with age has been observed as an increase in adolescence (5-20 years) (*19*), flattening in adulthood (20-40 years) (*19*) and a decline in old age (50-88 years) (*36*). Thus, the FHO model captures the age-related non-monotonic change in ΔPower (for the first time) naturally, while explaining HFD changes.

## Discussion

The present study uses a two-parameter model based on FHO to explain neural signals. We show that human life expectancy can be calculated using a single neural observable, i.e. the PSD slope rate. By far, this is the first study to determine lifespan, not the biological age, with a physiological parameter using a non-statistical model. Moreover, this is also the first model to our knowledge that captures (a) age-related slope and oscillations results simultaneously, (b) non-monotonic gamma power change with age naturally and (c) HFD change with age and stimulus.

### Neural mechanisms

The parameter *α* is inversely related to chronomemory, a measure of temporal history (distinct from cognitive memory). The age-related decline in *α* may reflect an accumulation of neural history or experience (*37*), thereby, leading to an increase in chronomemory. Conversely, the increase in *α* with stimulus may indicate a temporary suppression of chronomemory. During the stimulus presentation, the neural system prioritizes the immediate input over the preceding baseline activity, thus diminishing the influence of past activity and reducing chronomemory. As FD is inversely related to *α*, and therefore, directly to chronomemory, the increase in chronomemory may be related to asynchronous neural activity or decrease in long range temporal correlations, as suggested by FD interpretations (*9, 37*). Further, *α* may be a representative of multiple timescales involved in governing neuronal dynamics and adaptation (*38, 39*) or age-related long-term plasticity (*40*), that contributes to chronomemory.

The monotonic decrease of slope with age can help in constraining the neural processes governing it and subsequently *α*. The neural factors that had been proposed to modulate slope such as frequency filtering properties of dendrites and extracellular medium (*13*), tissue properties (*41*) and E-I balance (*15*), do not show a monotonic change from birth, indicating that they are not the main associate of human ageing. Even the variation of excitatory (Glutamate) and inhibitory (GABA) neurotransmitters with age is inconsistent in literature (*42, 43*). Interestingly, a metabolite, Taurine, which is involved in synaptic strength and neuronal excitability, had shown to decrease steadily in parietal and occipital grey matter from 0 to 18 years (*44*), further supported by other studies (*45, 46*). That may be the reason for good lifespan results in the present EEG dataset using the slope rate in the parietal and occipital electrodes. However, a recent study challenged Taurine as an aging biomarker using longitudinal data of blood samples (*47*), highlighting the complexity of identifying the neurochemicals controlling aging. Hence, the fundamental neural mechanism governing age-related changes in slopes, *α* and, thus, lifetime awaits to be discovered.

### Validity of assumptions

We constrain the model parameters *α* between 1 and 2 and *λ* between 0 and 1. In general, FHO is defined with order *α* varying between 0 and 2, with *α*_*FR*_ as *α* − *n* + 1; *n* being the greatest integer of *α*. As *α*_*FR*_ is same for *α* between (0,1] and (1,2], transitions between (*α* < 1 and *α* > 1 states lead to discontinuity in memory 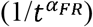. So, to avoid discontinuity, we have confined *α* between 1 and 2 only. The CF is given by 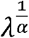. If *λ* is less than 1, CF decreases with decrease in *α*, and vice versa. With ageing, CF decreases from the observed trends while *α* decreases to explain slope results. So, to capture their concomitant decrease, we confine *λ* between 0 and 1.

We further assume *α* between 1 and 1.5 for baseline and 1.5 to 2 for stimulus-induced activity. The distinct *α* range for baseline and stimulus has enabled us to deal with slopes and oscillations independently and are equivalent of distinct neural mechanisms reflected in slopes (*13, 17*) and oscillations (*23, 24, 71*). Also, it explains the decrease of HFD during stimulus with respect to baseline. Moreover, it describes the observed non-monotonic variation of stimulus-induced gamma power with age (*19*). Narrowing the baseline range (e.g., 1 to 1.4) eliminates this trend, while extending the baseline regime shifts the gamma power peak from ∼20 years (Fig. 3) to 30–40 years, contradicting the observations (*19*).

Also, we assume the linear decrease of baseline *α* with age from 1.5 to 1. This assumption stems from the indirect assumption of the constant slope rate with age. While current literature lacks direct evidence to confirm or refute this, future studies may shed light on its validity.

### Factors affecting slopes

We could not directly map *α* to the measured slopes as we do not know the measured slopes close to birth and death and the theoretical slope differs from measured ones. The slopes in EEG and ECoG datasets lie between 0.5 to 2 (Fig. 2) while the theoretical slope from the model lies between 2 and 3 for 1 <*α* ≤ 1.5. The lower value of measured slopes could be attributed to white noise that may originate from recording system (*31, 72*). At frequencies higher than 100 Hz, the amplifier and thermal noise contribute primarily to electronic noise (*73*) and requires appropriate correction (*72, 74*). In addition, the slope measurement could be affected due to various factors, like reference scheme (*75*) and frequency range (*10*). Apart from external sources, the slopes could decrease due to spiking activity of neuronal population (*31, 76*). The present accuracy about lifespan through slope rate indicates that the noise is additive in nature and independent of age. Therefore, the change in parameters (here, slope) could lead to cancellation of such noise in the results. That was why, a strong correlation (-0.99) was observed previously between ‘changes’ in slope and HFD, rather than their absolute values (*9*). Similarly, here, the change of slope with ageing, the slope rate, not the slope itself, gives good lifespan estimation.

The slope rate is not always same in different conditions. For example, it was different between eyes open (0.010) and eyes closed (0.13) for healthy individuals as was reported in (*10*). We prefer eyes closed condition due to minimisation of EMG activity that could have aroused due to fixation in eyes open condition (*77*). We show that the suitable conditions with minimum noise exist, where the lifespan calculation from slope rate gives results in proximity with statistical lifespans. For example, both the datasets, used here, employed unipolar reference scheme and high frequency range (beyond 50 Hz) for slope computation. The slope change with age in EEG was found to prominent with unipolar reference (*7, 10–12*) and therefore, it may be the optimal reference scheme for lifetime calculations. Future standardization of factors can lead to improved accuracy of lifespan estimation.

### Implications in healthcare

The present mean lifespan estimation from a single and easy to measure variable could help in early risk detection of ‘lifespan-threatening’ diseases. A person with a shallower slope in high frequency range as compared to another age-matched person seems to have possibility of a lower healthy lifespan, thus, more possibilities of lifespan threatening diseases like Parkinson’s disease (*48, 49*) and schizophrenia (*50*). We do not find human slope studies at high frequencies (>50 Hz) for these diseases. Moreover, it could assist clinicians in drug testing and tumor detection possibilities and whether an individual has a normal lifespan after the treatment by comparing slope rate under these different conditions. Further, the correlation of slope rate with different physiological parameters like heart rate variability could result in determining early risks of non-neural organ diseases.

The high frequency slopes did not differ in patients suffering with mild cognitive impairment (MCI) and dementia due to Alzheimer’s disease (AD), as compared to their age and gender-matched healthy controls (*10*), suggesting its minimal role in reducing lifespan., This aligns with previous studies that showed dementia emerges only ∼0.8 years before death (*51, 52*). Given its typically late onset, AD-induced dementia seems to affect quality of life more than lifespan itself. Thus, this validates that slope at high frequencies can provide reliable measure of lifespan. Future experiments in mice and monkeys could help in further validation of this model.

## Conclusion

By providing average lifespan, the present model acts as the first step towards individualized lifetime predictions grounded in brain dynamics. It lays the foundation for understanding neural mechanisms as well as ageing through a lens of single paradigm that captures both linear and non-linear changes in neural signals. The model employed here could be instrumental in quantifying life using mathematics and unveiling its underlying fundamental principles. It holds promise for wide applications for theoretical and experimental neuroscientists, clinicians as well as policy makers, seeking objective metrics of ageing and longevity.

## Materials and Methods

### Model: Stochastic Fractional Harmonic oscillator (FHO)

We consider the stochastic fractional harmonic oscillator (FHO) of order *α* with 1<*α<* 2 (*54*). It is defined by

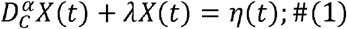

where 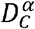 is the Caputo fractional derivative, *λ* is the decay constant, lying between 0 and 1 and *η* is the Gaussian normal noise. The Caputo derivative for *α* between 1 and 2 is represented as:

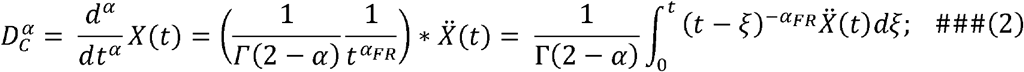

where * indicates convolution, 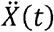 is the second order derivative of *X* (*t*) and *α*_FR_ is given by *α* − 1.

#### Solution

We compute the solution of equation (1) analytically using the following steps (*55, 56*):

I. **Taking Laplace Transform of the eq. (1):** Laplace Transform of *D* ^*α*^*X* (*t*) with *α* between 1 and 2 is

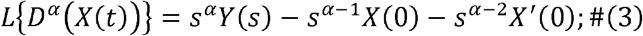

where *Y(s*) is the Laplace transform of *X(t*). Therefore, Laplace Transform of the Eq. (1) is

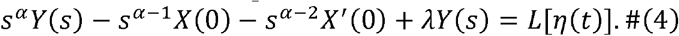

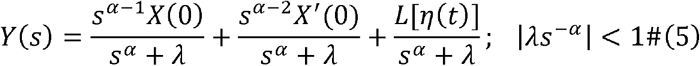

where *L* [*η(t*)] is the Laplace Transform of *η*(*t*).
II. **Taking Inverse Laplace Transform**

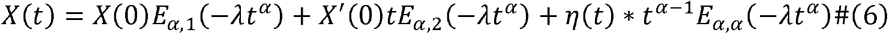

where the last term is the convolution of *η*(*t*) *and E* _*α,α*_ (− *λt* ^*α*^). *E*_*α, β*_(*z*) is the Mittag-Lefler function (*56*), defined by

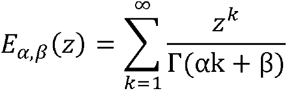

To characterize the power spectrum, we take the Fourier Transform *A*(*w*) of *X* (*t*) It can be obtained by replacing ‘*s’* by ‘*iw*’ and Laplace transform of Gaussian noise by its Fourier transform in Eq. (5).

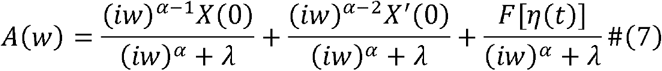

The Fourier transform of a Gaussian normal noise (with mean 0 and standard deviation 1) is a Gaussian normal noise in frequency domain. Thus,

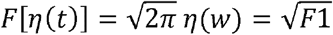

We assume that the system starts from rest. Therefore, we choose the initial conditions as *X(0) = 0* and *X’(0) = 0*. Hence,

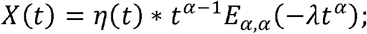

and

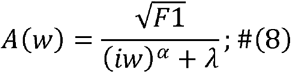

Therefore, power is given by

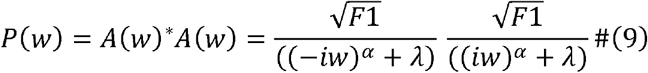

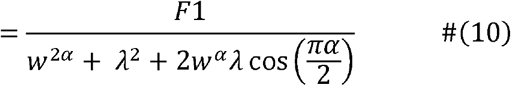

### Variation of Signal Features with *α* and *λ*

A. *Slope:* The aperiodic part of power (*P*_*AP*_ ; without peaks) (*53*) with offset ‘b’ and slope *χ*’ can be given by

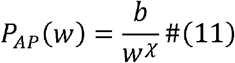 Comparing Eq. (11) with Eq. (10),

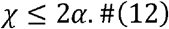 Thus, the slope increases with increase in *α*. Further, the equality in Eq. (12) holds if *λ* →0. As *λ* increases, slope decreases.
B. *Centre Frequency (CF):* The CF of oscillatory peak (for *α*> 1.5) or the knee frequency (for *α* < 1.5) can be computed using the diverging nature of Eq. (8). The denominator blows up at

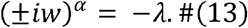 Thus,

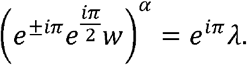 Therefore,

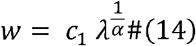

where *c*_1_ is 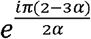 for + *iw* and 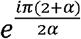 for − *iw*. Thus, *w* has complex roots. The CF can be computed as the real part of the above equation. Hence,

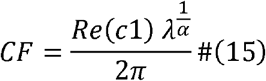 By Eq. (15), the CF increases with increase in *λ*. As *λ* < 1 and *α*> 1, increase in *α* leads to an increase in CF.
C. *Peak Height (PH):* To observe the variation of PH, we compute *P*(*w*) at absolute value of *w*, that represents the CF of peak using Eq. (15). So, substituting *w*∼*λ* ^1/*α*^ in Eq. (10), it becomes

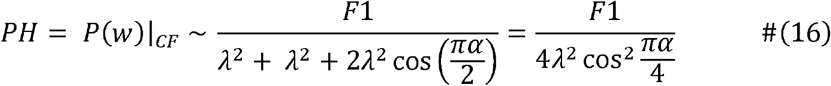 From the above expression, one can conclude that PH increase with decrease in *λ* and increase in *α*.
D. *Fractal Dimension (FD):* FD is a complexity measure that quantifies the irregularity and fractality of a signal (*57*). A higher FD indicates noise-like behaviour whereas smaller FD signifies smoothness and repeatability in signals. For a time series, it lies between 1 and 2; with 1 for a sinusoidal and 1<FD<2 for fractal signals and 2 for Gaussian noise, that fills the full space (*9, 58, 59*). Theoretically, it is related to a measure of long-range temporal correlations, the Hurst Index (H), as FD = 2 − H (*60, 61*). H characterizes the fractional Brownian motion (*58, 62*) and provides a basis for the anamolous and fractional diffusion, with the fractional order *α* varying as 2H (*63, 64*). By combining the dependence of FD and *α* on H, FD should be related to *α* as:

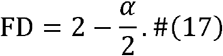

It implies that FD varies negatively with and does not depend on *λ*.

### Numerical Details

We obtain the time series using Eq. (6) with initial conditions as (0,0). The Mittag-Lefler function is computed using the matlab code from https://in.mathworks.com/matlabcentral/fileexchange/8738-mittag-leffler-function. We generate the signal at the sampling frequency of 2500 Hz for 0.5 s each, similar to the real EEG data used in (*9, 10*). We obtain 200 iterations for each condition of *α* and *λ*, varying *α* and *λ* in the steps of 0.05 between 1.1 and 1.95, and 0.1 between 0.1 and 1 respectively.

We first generate the signal in dimensionless units (*65*) for time *T*_*c*_ with the time step, *dt* _*c*_, as 0.1 (*38*). We, then, transformed the signal to the real time window *T*_*o*_ (in the present case, 0.5 s). The parameters *T*_*c*_ and *dt*_*c*_ could be set using the following relation between computational (dimensionless) and observational dimensional quantities, involving the number of total data points (N).

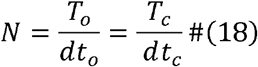

As the sampling frequency *F*_*s*_ is 1/*dt*_*o*_, therefore

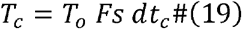

Thus, CF in observable dimensions (*CF*_*o*_), transformed from computational dimension (*CF*_*c*_) can be found using

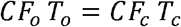

Hence

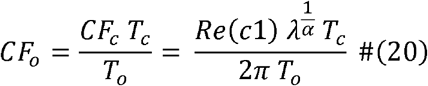

The second expression is obtained by replacing *CF*_*c*_ using Eq. (14). *α* can be determined using HFD ideally. Further, one can determine *λ* for the model appropriately using the oscillatory peaks, after the identification of *α*. It is important to note that for a given choice of peak frequency *λ* depends on the sampling frequency of the system as evident from Eqs. (19) and (20).

### Computation of CF, PH, slope and FD

We compute the PSD of the signal generated using Eq. (6) for each trial (repeat) using Chronux toolbox (((*66*); RRID:SCR_005547) via multi-taper analysis with single taper. We then obtain the trial-averaged PSD. We define the CF and PH for an oscillatory peak as the frequency and the corresponding power at which the trial-averaged PSD is maximum. We use Matlab wrapper for Fitting Oscillations and One Over F (FOOOF) toolbox to compute the slope (*53*), in frequency range of 64-140 Hz similar to the low frequency range of (*10*). We choose the parameters as aperiodic mode: ‘fixed’, peak threshold: 2.0, maximum number of peaks: 1 and peak width limits: [4 80]. We compute FD, using Higuchi’s method (*67*), the Higuchi fractal dimension (HFD) for each trial using a custom Matlab code and set *k*_*max*_ 10 similar to (*9*).

### Neural Data analysis

#### EEG analysis

To compute the slope rate with respect to age, we take resting state eyes closed EEG data. We preferred the eyes closed condition over eyes open to avoid artifacts due to electromyography (EMG) activity. We use the clean eyes closed data of 204 healthy subjects aged between 50-88 years. The data were collected under Tata Longitudinal Study of Aging (TLSA) (*8, 10*). The details of experimental setup, data collection and artefact rejection for eyes closed condition could be found in (*10*). Here, we provide their summary.

The TLSA EEG dataset for eyes closed condition was collected from 223 cognitively healthy humans aged 50-88 years from the urban community of Bengaluru, Karnataka with the informed consent and monetary compensation. The resting state eyes closed EEG data were recorded using 64-channel active electrodes (actiCap) from BrainAmp DC EEG acquisition system (Brain Products GmbH) for 1-2 minutes in a dark room. The electrodes were placed following international 10–10 system and were referenced online at FCz. EEG signals were sampled at 2.5 kHz. They were filtered online between 0.016 Hz (first-order filter) and 1 kHz (fifth-order Butterworth filter) and had 16-bit resolution (0.1 μV/bit). The data were segmented into 2s non-overlapping epochs/ trials, resulting a resolution of 0.5 Hz in frequency domain.

The topoplot function of EEGLAB toolbox ((*68*), RRID:SCR_007292) was used to generate scalp maps with standard *Acticap 64* unipolar montage. In the previous studies (*8–10, 36, 69*), main analyses were done on a subset of occipital and centro-parietal groups comprising P3, P1, P2, PO3, POz, PO4, O1, Oz and O2, as strong gamma power (power in 20-35 Hz) was observed for them. They were termed as ‘high-priority’ electrodes. We used the analysis from the same set of electrodes.

To obtain clean data, an artifact rejection pipeline had been used. It involved (i) the rejection of electrodes with impedance greater than 25 kΩ; (ii) the detection of bad epochs as the epochs with root mean square (RMS) value outside 2.5-35 *μ* V and the deviation more than 6 standard deviation in frequency domain; (iii) the removal of electrodes with more than 30% bad epochs; (iv) rejection of epochs that were bad in the high priority electrodes or in more than 10% of other electrodes; and (v) the rejection of electrodes whose slopes for power spectrum between 56 and 84 Hz were less than 0. Overall, it led to the rejection of 3.11 ± 4.87% (mean ± SD) epochs, 18.86 ±13.10% electrodes and 10 subjects. Additionally, 9 participants were rejected who were recorded using 32-channel EEG system. It resulted in the clean eyes closed data for 204 subjects (112 males and 92 females) with average 39.35 ±7.75 epochs for each subject. The unipolar reference scheme was used for analysis.

The PSD was computed using the multi-taper method with a single taper, implemented via the Chronux Toolbox ((Bokil et al., 2010); RRID:SCR_005547). Spectra were first estimated for individual epochs and then averaged across epochs for each electrode. Slopes were computed for the epoch-averaged PSD between 64-140 Hz using FOOOF toolbox (*53*).

#### Electrocorticography (ECoG) analysis

We use the ECoG data of (*7*) for the computation of slope rate. As the data was not readily available online, we obtain the slope for different ages in frequency range of 75-150 Hz from Fig. 2B of (*7*). Briefly, the ECoG data were recorded from 15 epileptic patients (11 male; 4 female), aged 15-53 years, using chronic subdural electrodes with informed consent. The subjects performed auditory tasks that included passive phoneme listening, word repetition or auditory attention tasks. The data were analysed from frontal, temporal and supramarginal neocortical regions after amplification X10,000, analog filtering (0.01–1000 Hz), digitization, and downsampling to 1000 Hz. The average reference scheme was implemented and the data were high pass filtered above 1 Hz using finite impulse response (FIR) filter. Epileptic electrodes and epochs and electrodes with low signal-to-noise-ratio were discarded from analysis. The PSDs and slopes were computed using Welch’s method for 2 s time window with 50% overlap and linear regression respectively.

#### Slope rate calculation

To calculate the slope rate, we perform linear regression between slope vs age. We use *fitlm* function in Matlab that generates the slope rate and p-value using t-statistics.

### Statistical analysis

The correlation coefficients (*ρ*) and the corresponding p-value for HFD vs slope and CF vs age are obtained using the Matlab command *corrcoef*. For linear regression between different quantities, we use *fitlm* function in Matlab that generates the model parameters *β,y*0 corresponding to *y= y*0 + *β *x* and p-value using t-statistics.

## Acknowledgments

We are grateful to Prof. Banibrata Mukhopadhyay, Department of Physics, IISc for discussion and support in framing and writing this work. We thank Prof. Supratim Ray, Centre for Neuroscience, IISc for providing TLSAEEG eyes closed data as well as feedback and suggestions that enhance the quality of the present work. We also thank Dr. Surya Prakash for scientific discussions. We use adapted versions of Fig. 6 bottom left subplot of (*10*) under Creative Commons CC BY license and Fig. 2B of (*7*) under Creative Commons Attribution 4.0 International License (CC-BY).

## Data and Code availability

Codes to generate the simulated data and figures are available on GitHub at https://github.com/4srihy/fractionalDynamicsNeuralModel. All the modelling and data analyses were done using Matlab R2021b. The Mittag-Lefler function was computed using the Matlab code available at https://in.mathworks.com/matlabcentral/fileexchange/8738-mittag-leffler-function.

The PSDs were obtained using Chronux toolbox (version 2.10), available at http://chronux.org. Slopes were computed using matlab wrapper for FOOOF (*53*) (https://github.com/fooof-tools/fooof_mat). HFD was calculated using custom code based on https://in.mathworks.com/matlabcentral/fileexchange/124331-higfracdim.

## Supplementary Information

Materials and Methods

Fig. S1

Supplementary Text

1. Simultaneous occurrence of peak and slope in PSD of neural signal (includes Fig. S2)
2. Slope rate analysis for gender and electrode position from TLSA EEG dataset (includes Fig. S3)
3. Stochastic Damped Harmonic Oscillator (includes Fig. S4)

## Supplementary Information

### Supplementary Figure

**Figure S1.**
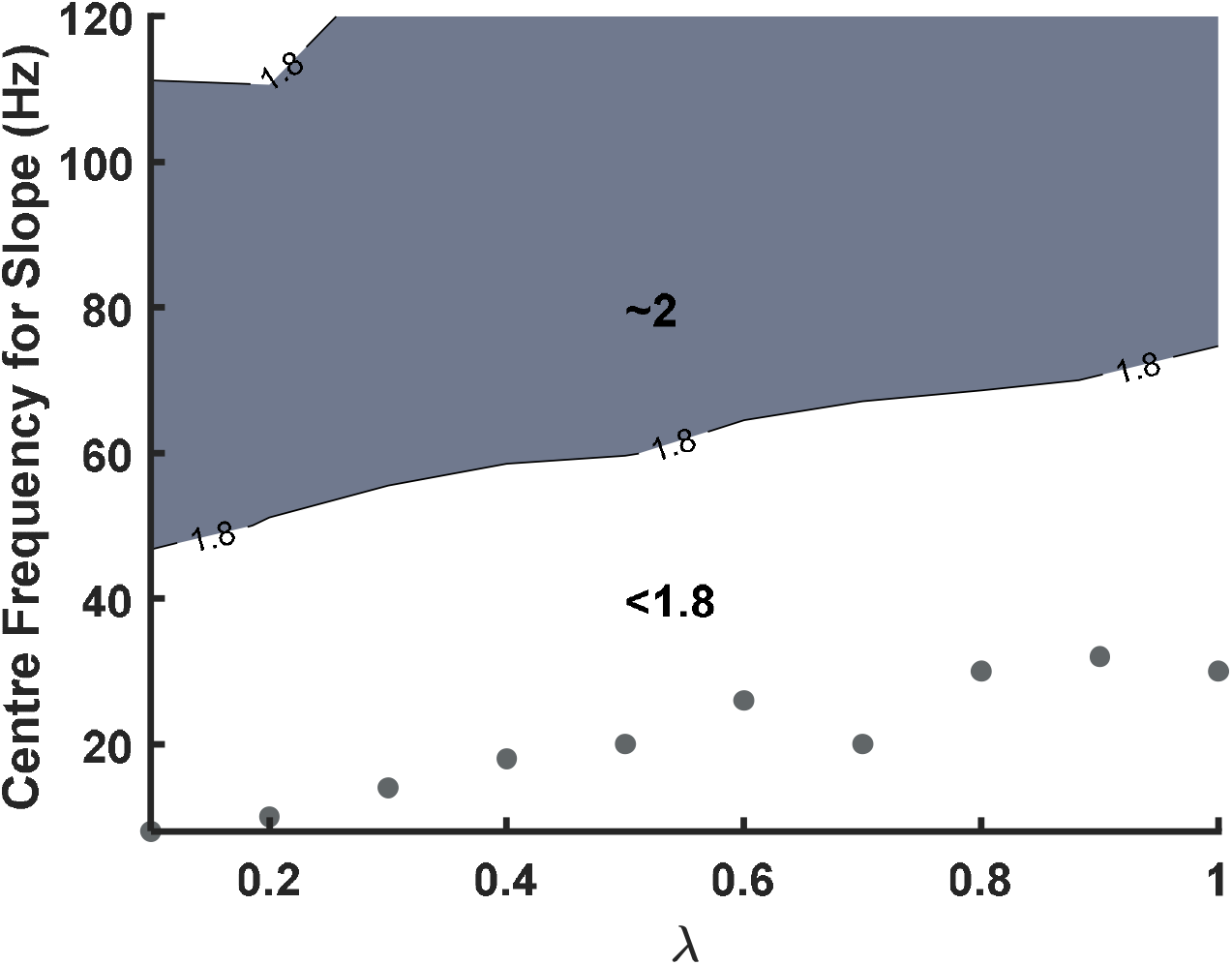
Variation of dSlope/d*α* with *λ* and centre frequency. *dSlope/dα* has been calculated using the slope values for *α* between 1.1 and 1.5. The slope for each *α* is computed for the frequency ranges of frequency width 80 Hz between 4 to 160 Hz with centre frequency starting from 20 Hz in steps of 10 Hz. The frequency ranges with centre frequency 20 and 30 Hz have lower frequency width due to our restriction of lower cut-off as 4 Hz. The grey region shows *dSlope*/*dα* ∼2 (between 1.8 and 2.2) and the white region displays *dSlope*/*dα*<1.8. The grey dots correspond to the knee frequency at a given *λ* at *α* = 1.5. The condition *dSlope*/*dα* ∼2 is attained only at high frequencies, particularly away from knee, suggesting that age estimation should be done using slopes at high centre frequencies.

## Supplementary Sections

### Section 1. Simultaneous occurrence of peak and slope in PSD of neural signal

The resting state activity in EEG is not completely devoid of peaks. Instead, there may be low frequency peaks representing oscillation in alpha (8-12 Hz) or beta (15-30 Hz) range, along with slope. In such cases, the PSD of neural signal can be computed as the sum of two distinct PSDs corresponding to different *α*: one representing the aperiodic slope component (*α*_*slope*_ < 1.5) and the other for oscillatory peak (*α*_*osc*_ > 1.5). This approach is aligned with the technique used by Donoghue and colleagues in the FOOOF framework (*53*), where they had separated peaks and slope in PSD before their characterization. As an example, we show the PSDs for *α* = 1.2 and 1.8 at *λ* = 0.2 for slope and peak characterization respectively in Fig. S2. The PSD in the dotted line is the sum of PSDs obtained from these two *α*. It is characterized by slope of PSD with *α* = 1.2 and peak of PSD with *α* = 1.8. Though there is no change in PH and CF of oscillatory peak in the resultant PSD, it has a higher slope of 2.73 in the frequency range of 64-140 Hz as compared to the slope of 2.32 for *α* = 1.2. Therefore, slopes computed in the vicinity of peaks may be confounded as was also discussed in (*31*). Moreover, if peak and slope characteristics are used from a single simulated PSD, it would imply that they are governed by a single neural mechanism and should follow a strong correlation, which is not always the case.

**Figure S2.**
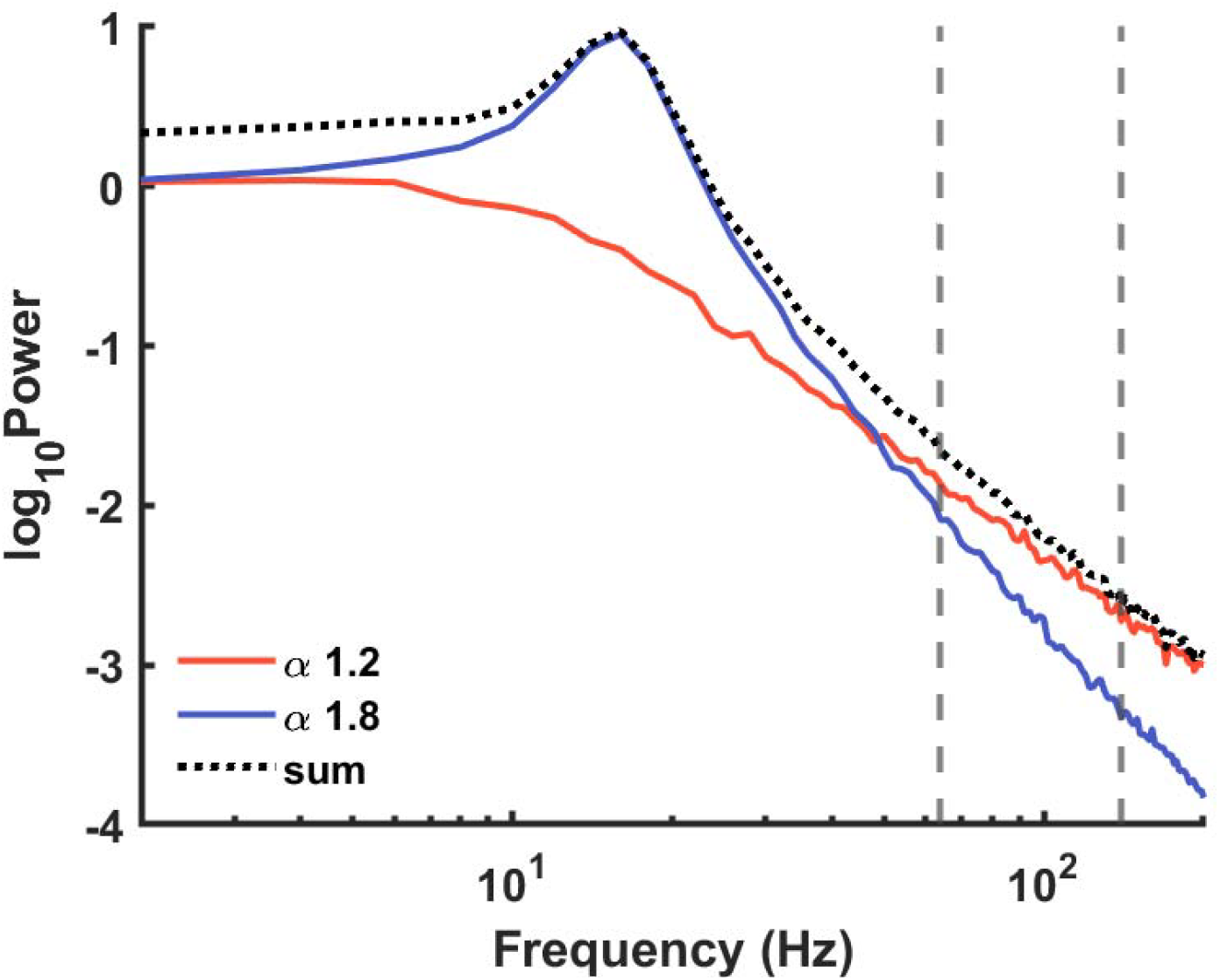
The resultant PSD for peak and slope characterization. Red and blue (solid) lines represent trial-averaged PSD for *α* 1.2 and 1.8 respectively for *λ* 0.2. The sum of these PSDs are shown as black dotted line. The grey lines mark the region of slope computation (64-140 Hz).

### Section 2: Slope rate analysis for gender and electrode position from TLSA EEG dataset

We calculate the slope rates for each gender using TLSA EEG eyes closed dataset of healthy individuals (*10*). For the group of occipital and parietal electrodes, the slope rates for males (N = 112) and females (N = 92) are −0.011 ± 0.0006 (p = 9.21 ×10^−4^ and −0.013 ± 0.0009 (p = 0.0045) yielding the maximum life expectancy of 90.9 [86.2, 96.2] years and 76.9 [71.9, 82.6] years respectively. These results are opposite to the population statistics having life expectancy of female (71.3 years) higher than males (67.9 years; (*32*)). It could be due to lesser number of subjects in the gender groups (∽100 each) as compared to the total subjects (204) resulting in a less reliable slope rate measurement for each gender.

Next, we test whether slope rate depends on electrode position. We compute slope rate for each electrode. The topoplot in Fig. S3 shows the variation of slope rate across whole head map. The red dots indicate the electrodes for which slope rate was significant (p<0.05, *t-test*, false discovery rate (FDR) corrected). The slope rate across the 40 significant electrodes is −0.0133 ± 0.0001 (mean ± SEM), giving a similar maximum life expectancy of 75.19 [74.63,75.76] years, as reported in the main text. Thus, (significant) slope rates do not vary much with electrode position, except the frontal electrodes that have slope rate < − 0.02.

**Figure S3.**
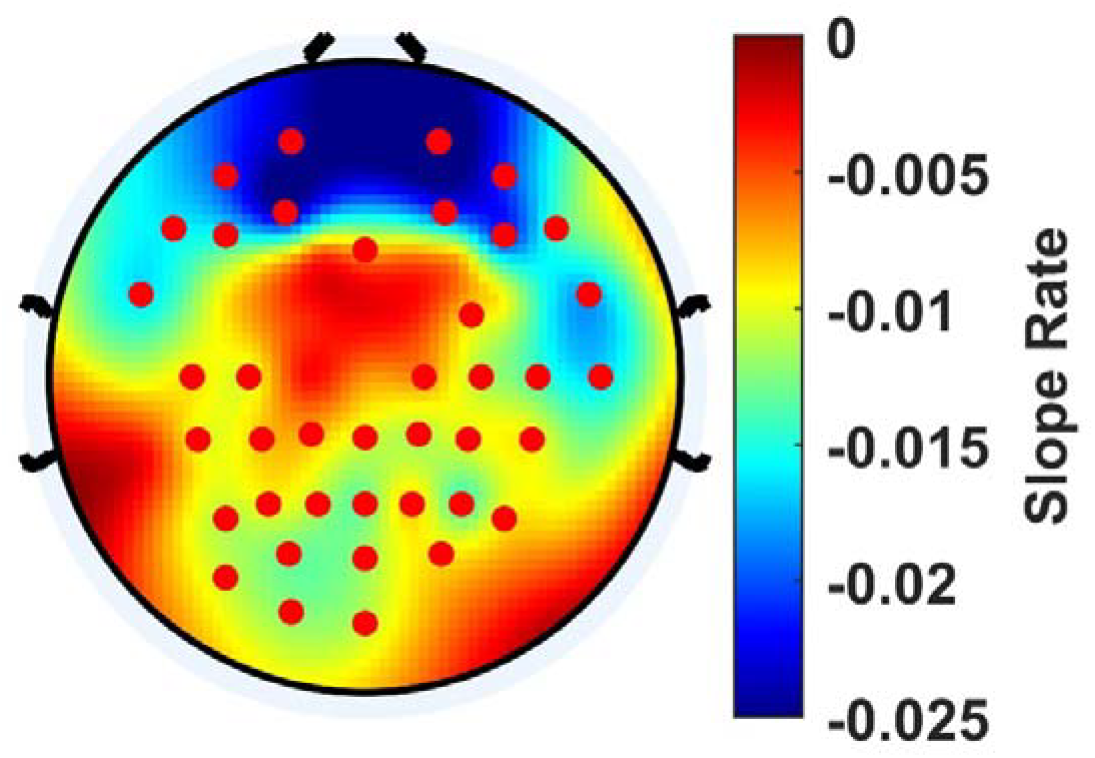
Topoplot showing the variation of slope rate, computed for frequency range 64-140 Hz for all 64 electrodes for eyes closed data from TLSA EEG dataset. The electrodes highlighted in red have significant slope rate with p<0.05, *t-test*, false discovery rate (FDR) corrected.

### Section 3. Stochastic Damped Harmonic Oscillator

We compared our model of FHO with auto regressive model with 2 parameters (AR(2)), which corresponds to a stochastic damped harmonic oscillator (DHO) (*70*). Briefly, the time points *x*_*t*_ in an AR(2) model are described by

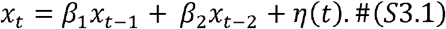

The above equation can be simplified using lag operator form as

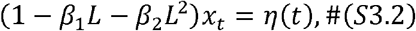

where *L* is the lag operator such that *Lx*_*t*_ = *x*_*t* − 1_. We can obtain the eigenvalues of the above equation as the roots (*m*) of the characteristic polynomial

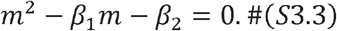

This is the eigenvalue equation for stochastic damped harmonic oscillator described by Eq. (S3.4) with *β*_1_ as the damping factor *b* and *β*_2_ as 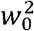.

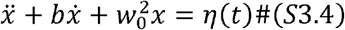

The eigenvalues are

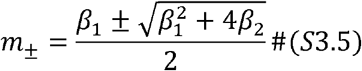

The DHO shows the oscillatory behaviour with decaying amplitude if the eigenvalues are complex conjugates with |*m* _±_ |<1. From Eq. (S3.5), these conditions are satisfied if |*β*_1_| < 2 and 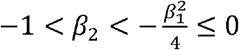.

We solve the Eq. (S3.1) using custom Matlab code. We choose the parameters, *β*_1_ between 1.8 and 1.99, *β*_2_ between -1 and -0.9890 and the initial conditions as the resting state with (*x*_1_, *x*_2_) as (0,0). The variation of the PSD features and HFD is shown in Fig. S4. While FHO shows both noise like and oscillatory behaviour (at different *α*), DHO displays only oscillatory behaviour as can be observed in time series with the resemblance with smooth sinusoids as well in PSDs that always have oscillatory peaks (Fig. S4 A and B). The slopes are dependent on PH and follow an inverse relation with PH for both *β*_1_ and *β*_2_ (Fig. S4C). Therefore, DHO cannot explain the simultaneous decrease in baseline slope (*10*) and oscillatory PH (*8*) with age. Further, both HFD and slope vary inversely with *β*_1_ (Fig.S4C), thus, varying directly with each other. This is in contrast with previous studies showing the inverse relation of FD with slope (*9, 67*). The variation of PH with HFD is parameter dependent. It varies negatively due to *β*_1_ but varies in the same direction as HFD due to *β*_2_ (Fig.S4D). However, for FHO, the PH varies inversely with HFD due to both *α* and *λ* (Fig. 1), similar to the previously observed variation (*9*). Overall, DHO fails to explain age-related slope, power and HFD results simultaneously.

A similar issue has been reported by a recent study (*22*), that tries to explain age-related slope and oscillatory power results in resting state using compartmental model. Though, it explained age-based decline in slope and CF, but failed to capture the power changes, since compartmental model is a linear model like DHO.

**Figure S4.**
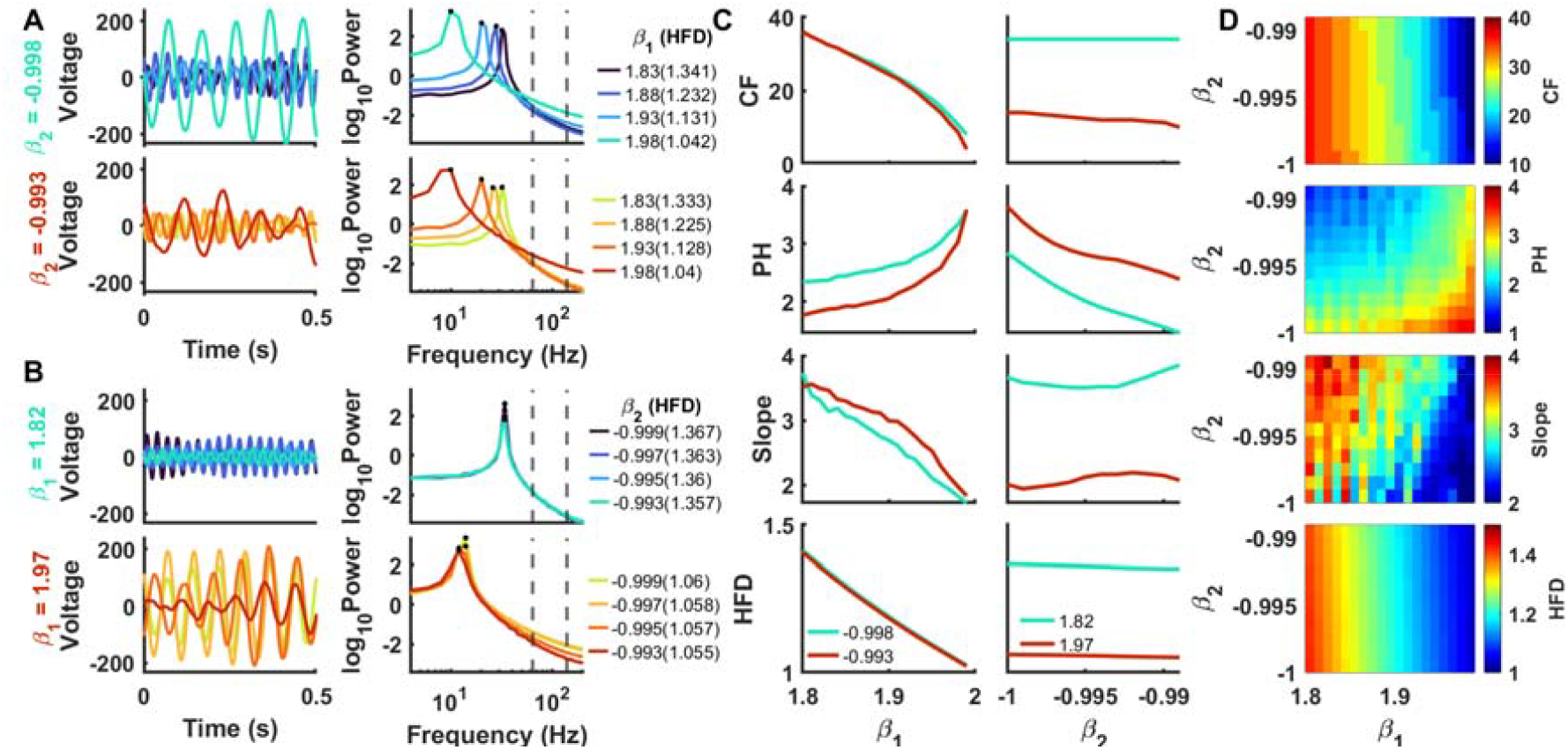
Variation of signal features with and for DHO. Same as Fig. 1 but for DHO.

